# An intestinal sphingolipid promotes neuronal health across generations

**DOI:** 10.1101/2022.07.24.501274

**Authors:** Wenyue Wang, Tessa Sherry, Xinran Cheng, Qi Fan, Rebecca Cornell, Jie Liu, Zhicheng Xiao, Roger Pocock

## Abstract

Maternal diet and environment can influence the neuronal health of offspring. Here, we report that diet-induced intestinal sphingolipid biosynthesis reduces adult-onset neurodegeneration intergenerationally in *Caenorhabditis elegans*. Feeding *C. elegans* with ursolic acid (UA), a natural plant product, provides neuroprotection by enhancing maternal provisioning of sphingosine-1-phosphate (S1P) - a bioactive sphingolipid. S1P promotes neuronal health across generations by upregulating transcription of the acid ceramidase-1 (*asah-1*) gene in the intestine. Intergenerational intestine-to-oocyte S1P transfer is essential for promoting neuronal health and is dependent on the lipoprotein yolk receptor RME-2 (Receptor-Mediated Endocytosis-2). Spatially regulating sphingolipid biosynthesis is critical, as inappropriate *asah-1* neuronal expression causes developmental axon outgrowth defects. Our results reveal that sphingolipid homeostasis impacts the development and intergenerational health of the nervous system.

**One-Sentence Summary:** An intestinal lipid prevents neurodegeneration across generations.

## Main Text

Axons are long cytoplasmic projections that transmit information between neurons. Maintenance of neuronal health requires the transport of cargo (organelles, RNAs, proteins and lipids) along the axonal cytoskeleton (*1, 2*). Microtubules are major cytoskeletal components comprised of cylindrical structures assembled from α- and β-tubulin heterodimers that are crucial for intracellular transport (*1, 3*). Defective microtubule structure disrupts the supply of essential materials and is associated with multiple neurodegenerative disorders (*4*). Therefore, it is crucial to identify molecular mechanisms that support axonal health under conditions of sub-optimal microtubule-associated intracellular transport.

We used the *Caenorhabditis elegans* mechanosensory neurons to identify molecules important for maintaining the structural integrity of axons. The posterior lateral mechanosensory (PLM) neurons extend axons along the length of the worm to coordinate touch responses (*5*). A previous study showed that loss of the MEC-17/αTAT1 α-tubulin acetyltransferase causes adult-onset and progressive PLM axon degeneration (*6*). MEC-17 loss causes microtubule instability and aberrant axonal transport, leading to reduced number and disrupted spatial distribution of mitochondria, as well as defective synaptic protein localization (*6*). Neurodegeneration is caused by axon fragility, as the phenotype is robustly suppressed by paralyzing *mec-17* mutant animals (*6*). Further, PLM axon degeneration in *mec-17* mutants is exacerbated in animals with increased body length (e.g. *lon-2* mutant), likely due to the added demand on axonal transport required to maintain a longer axon (*6*).

To identify molecules that suppress axon degeneration, we expressed green fluorescent protein (GFP) specifically in the *C. elegans* mechanosensory neurons of *mec-17(ok2109); lon-2(e678)* mutant animals (Fig. 1). In a screen of natural products, we identified ursolic acid (UA) as a suppressor of axon degeneration in *mec-17(ok2109); lon-2(e678)* mutant animals (Fig. 1). UA is a lipophilic pentacyclic triterpenoid acid found in plants that has broad biological functions, acting as an anti-inflammatory, antioxidant, and neuroprotective molecule (*7, 8*). To assess the potency of UA-induced neuroprotection, we fed *mec-17(ok2109); lon-2(e678)* animals with different UA concentrations and examined PLM axon degeneration in day 3 adults (Fig. 1A-D and fig. S1A). We found that incubating P0 larval stage 4 (L4) animals with 50 μM UA for one generation robustly reduced axon degeneration in *mec-17(ok2109); lon-2(e678)* F1 progeny (Fig. 1C-D). To determine if the UA-induced suppression of axon degeneration was caused by reduced body length or motility we used Wormlab tracking (MBF Bioscience LLC, Williston, VT USA). We found no change in motility or body length of *mec-17(ok2109); lon-2(e678)*animals exposed to UA compared to controls, suggesting a molecular rather than physical effect (fig. S1B-E).

**Fig. 1.**
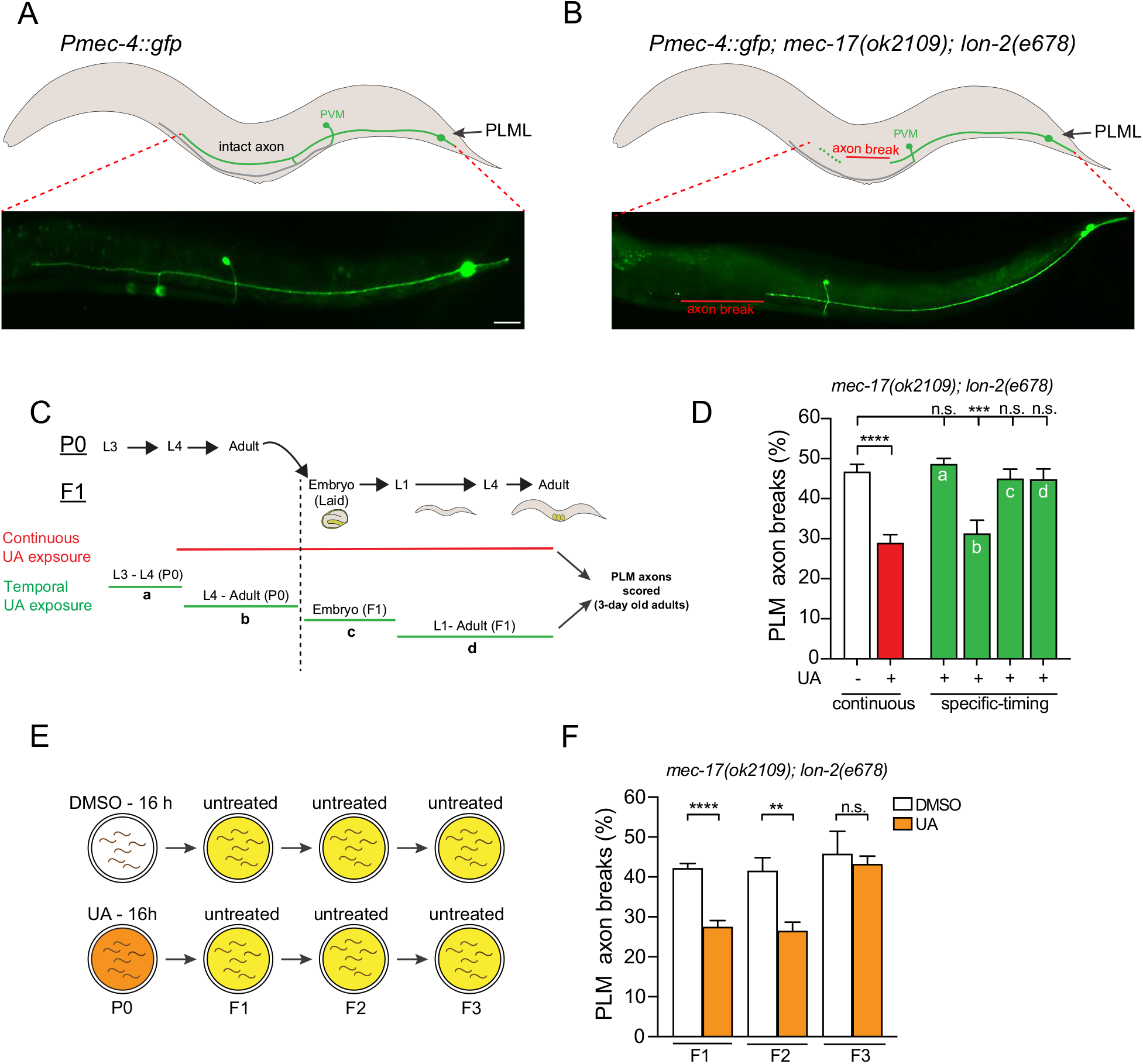
Ursolic acid promotes neuronal health across generations. (**A** and **B**) Schematics (top) and fluorescent micrographs (bottom) of PLM anatomy in wild-type (**A**) and *mec-17(ok2109); lon-2(e678*) (**B**) animals expressing the *Pmec-4::gfp* transgene (*zdls5*). A typical PLM axon break (red line) observed in 3-day old adult *mec-17(ok2109); lon-2(e678)* animals is indicated (**B**). Left lateral view, anterior to the left. Scale bar: 25 μm. (**C**) Ursolic acid (UA) exposure timeline. Upper schematic shows the stages of *C. elegans* development relevant to UA exposure in the P0 and F1 generations. Red line = continuous UA exposure from P0 L4 to F1 3-day old adult; green lines = specific timing of UA exposure. Vertical dashed line = demarcation between P0 and F1 generations. (**D**) Continuous exposure (P0 - L4 larva to F1 - adult) of UA rescues PLM axon breaks in *mec-17(ok2109); lon-2(e678)* animals (red bar). UA exposure of P0s from L4 larva to adult (b) but not earlier (a) or later stages (c and d) rescues PLM axon breaks in *mec-17(ok2109); lon-2(e678)* animals. n = 107 to 249. *P* values from one-way analysis of variance (ANOVA). \*\*\**P* ≤ 0.001; \*\*\*\**P* ≤ 0.0001; n.s., not significant. Error bars indicate SEM. (**E**) Experimental scheme for intergenerational inheritance experiment. P0s (L4 larvae) were treated with DMSO (control) or UA for 16 h. Eggs were laid on untreated plates for 3 h and resultant L4 larvae transferred to untreated plates for subsequent generations. (**F**) UA exposure in P0 mothers for 16 h (L4 - young adult) rescues PLM axon breaks in *mec-17(ok2109); lon-2(e678)* animals for two generations (F1 and F2). n = 77 to 128. *P* values from one-way analysis of variance (ANOVA). \*\**P* ≤ 0.01; \*\*\*\**P* ≤ 0.0001; n.s., not significant. Error bars indicate SEM.

We wondered whether there was a critical functional period during development for UA to reduce PLM axon degeneration. To investigate this, we exposed *mec-17(ok2109); lon-2(e678)* animals to UA at the following stages of *C. elegans* development: P0 L3 larvae-L4 larvae (during sperm generation and before oocyte production), P0 L4 larvae-adult (oocytes and sperm present), embryo-L1 larvae (embryogenesis) or L1 larvae-adult (all larval stages and adult). We found that axon degeneration in F1 adult progeny was only reduced when P0 hermaphrodites containing oocytes and sperm (P0 L4-adult) were exposed to UA (Fig. 1C-D). This result suggested that the axonal health-promoting effect of UA is deposited in the gametes to maintain axonal health through to adulthood. This prompted us to examine if UA could also protect the nervous system in subsequent generations. Incubating P0 animals with UA from L4 larvae-adult (16 hours) reduced axonal degeneration for two, but not three, generations - revealing an epigenetic intergenerational effect (Fig. 1E-F).

How does UA intergenerationally protect the nervous system? As the UA functional period is during oocyte production (P0 L4 larvae-adult), we hypothesized that UA may affect maternal yolk provisioning to oocytes. Yolk is synthesized in the hermaphrodite intestine and contains lipids and lipoproteins that provide oocytes, and thus embryos, with nutrients for development (*9*). Oocyte yolk import occurs through endocytosis and requires RME-2 (Receptor-Mediated Endocytosis-2), a low-density lipoprotein receptor (*10*). We found that RME-2 is required for UA to reduce PLM neuron degeneration, supporting a role for intestine-oocyte transport (Fig. 2A). Animals lacking RME-2 are also defective in transport of RNAs that are major transmitters of epigenetic inheritance (*11*). The nuclear HRDE-1 (Heritable RNAi Deficient 1) Argonaute is required for inheritance of small RNAs, however, we found that *hrde-1* is dispensable for UA to reduce PLM degeneration (fig. S2) (*12*). These data suggest that UA induces alternative epigenetic factors in the maternal yolk and that this information optimizes the oocyte/embryonic environment to promote neuronal health.

**Fig. 2.**
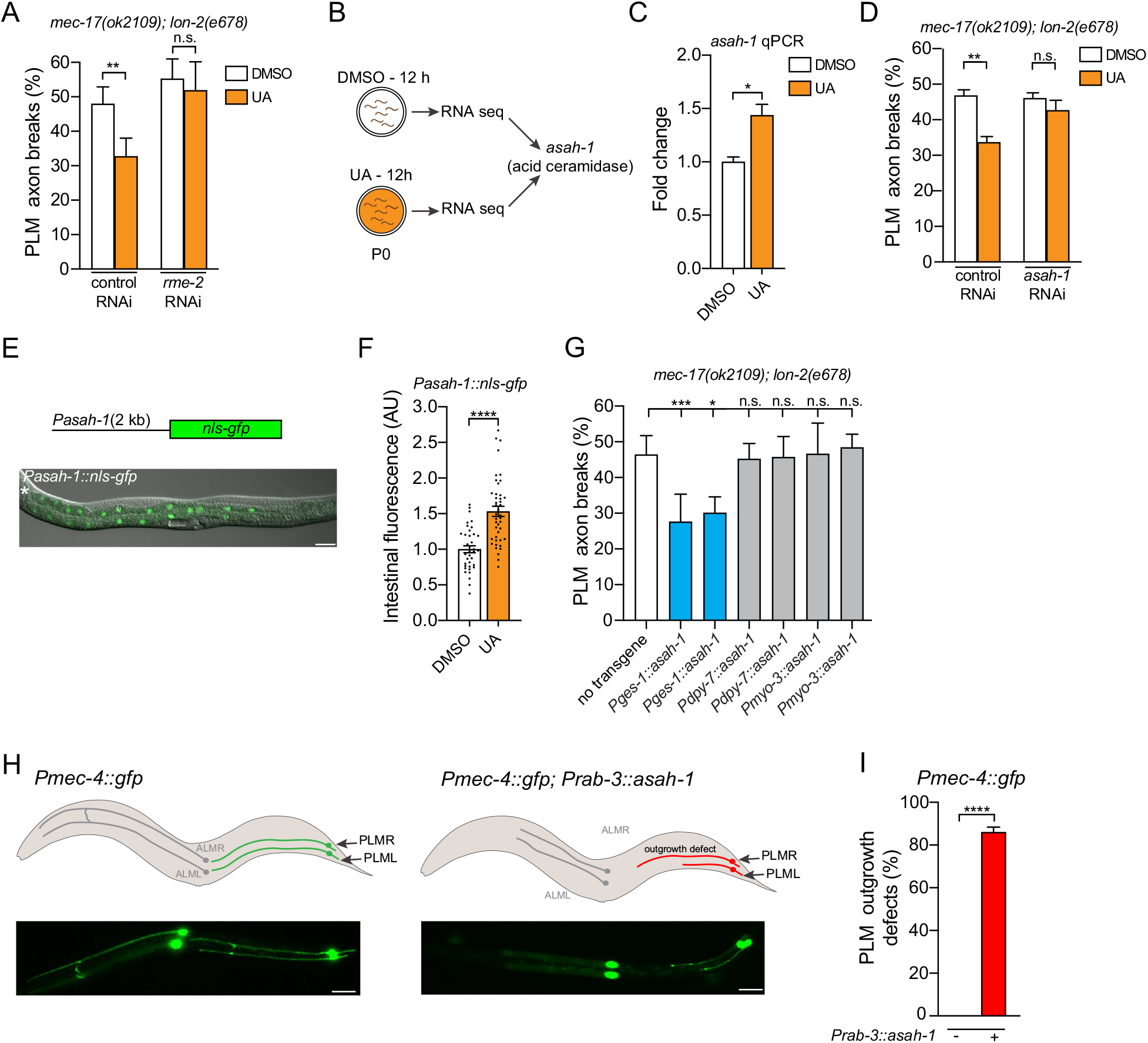
Ursolic acid induces acid ceramidase (*asah-1*) expression to promote neuronal health. (**A**) *rme-2* RNAi knockdown suppresses the ability of UA to rescue PLM axon breaks in *mec-17(ok2109); lon-2(e678)* animals. n = 154 to 187. *P* values from one-way analysis of variance (ANOVA). \*\**P* ≤ 0.01; n.s., not significant. Error bars indicate SEM. (**B**) Schematic of transcriptome analysis following UA exposure. Total RNA isolated from L4 larvae exposed to DMSO or UA for 12 h. *asah-1* is upregulated following UA exposure (Table S1). (**C**) *asah-1* mRNA is upregulated in wild-type L4 larvae after 12 h of UA exposure. *P* values assessed by unpaired t test. \**P* ≤ 0.05. Error bars indicate SEM. Triplicate biological replicates. *cdc-42* housekeeping gene used as a control. (**D**) *asah-1* RNAi knockdown suppresses the ability of UA to rescue PLM axon breaks in *mec-17(ok2109); lon-2(e678)* animals. n = 94 to 148. *P* values from one-way analysis of variance (ANOVA). \*\**P* ≤ 0.01; n.s., not significant. Error bars indicate SEM. (**E** and **F**) A *Pasah-1::nls-gfp* transcriptional reporter (*rpIs165*) drives expression in the intestine from late embryogenesis through to adult (see fig. S4) - L4 larva shown in (**E**). *Pasah-1::nls::gfp* expression increases in animals exposed to UA for 12 h compared to control (DMSO) (**F**). Lateral view, anterior to the left. Scale bar: 25 μm. n = 37 to 43. *P* values assessed by unpaired t test. \*\*\*\**P* ≤ 0.0001. Error bars indicate SEM. (**G**) Driving *asah-1* cDNA expression using the heterologous intestinal promoter (*ges-1*) but not the hypodermal (*dpy-7*) or muscle (*myo-3*) promoters rescues PLM axon breaks in *mec-17(ok2109); lon-2(e678)* animals. n = 72 to 305. Error bars indicate SEM. *P* values from oneway analysis of variance (ANOVA). \**P* ≤ 0.05, \*\*\**P* ≤ 0.001. n.s., not significant. (**H** and **I**) Overexpression of *asah-1* in the nervous system (*rab-3* promoter) causes developmental axon outgrowth defects in mechanosensory neurons of wild-type animals expressing the *Pmec-4::gfp* transgene (*zdIs5*) Schematics and fluorescent micrographs of ALM and PLM axons in wild-type (left) and *Prab-3::asah-1* expressing animals (right) (**H**). Quantification of PLM axon outgrowth defects of *Pmec-4::gfp* and *Pmec-4::gfp; Prab-3::asah-1* expressing L1 larvae (**I**). *P* values assessed by unpaired t test. *\*\*\*\*P* ≤ 0.0001. n = 62 to 95. Error bars indicate SEM. Typical axon outgrowth defect observed is marked in red (**H**). Left lateral view, anterior to the left. Scale bar: 25 μm.

To identify factor(s) regulated by UA to prevent PLM axon degeneration, we examined the transcriptomes of synchronized L4 larvae that were exposed to UA (or DMSO as a control) for 12 hours (Fig. 2B, fig. S3 and Table S1). We identified 49 dysregulated genes (FDR 0.02) (8 upregulated and 41 downregulated) and surveyed this dataset for genes expressed in the intestine (the source of yolk) that potentially control lipid metabolism. We detected increased *asah-1* transcript abundance in animals exposed to UA, which we confirmed by quantitative polymerase chain reaction (qPCR) analysis of independent RNA samples (Fig. 2C). *asah-1* encodes an acid ceramidase that hydrolyzes ceramide into fatty acid and sphingosine (*13*). To assess whether *asah-1* is required for UA to reduce PLM axon degeneration we performed RNA-mediated interference (RNAi) to knock down *asah-1* in *mec-17(ok2109); lon-2(e678)* animals incubated with UA (Fig. 2D). We found that UA does not reduce PLM axon degeneration in *asah-1* RNAi knockdown animals, suggesting that *asah-1* expression is required for the UA neuroprotective effect (Fig. 2D). To identify the tissue in which *asah-1* is expressed and potentially regulated by UA, we monitored the spatial and temporal expression pattern of *asah-1* using 2000 bp of upstream sequence to drive nuclear-localized expression of the gene encoding green fluorescent protein (*gfp*) (Fig. 2E and fig. S4). We detected nuclear GFP expression exclusively in intestinal cells from late embryos through to adult in this *Pasah-1::gfp* transcriptional reporter (Fig. 2E and fig. S4). At all stages, we observed a bias in nuclear GFP expression in the anterior intestine (fig. S4). We imaged nuclear GFP expression in *Pasah-1::gfp* L4 animals after 12 h of UA exposure and detected increased fluorescence in intestinal nuclei (Fig. 2F). Thus, UA induces *asah-1* transcription in the intestine.

As *asah-1* is upregulated in animals exposed to UA, we wondered whether transgenic overexpression of *asah-1* in *mec-17(ok2109); lon-2(e678)* animals could mimic the neuroprotective effect of UA. Single cell sequencing studies corroborate that *asah-1* is predominantly expressed in the intestine, however low-level expression is detected other cells/tissues, including the PLM neurons (*14, 15*). We therefore overexpressed *asah-1* in *mec-17(ok2109); lon-2(e678)* animals using the following heterologous promoters: intestine (*ges-1* promoter), hypodermis (*dpy-7* promoter), muscle (*myo-3* promoter) and the mechanosensory neurons (*mec-4* promoter) (Fig. 2G) (*16–19*). Overexpressing *asah-1* in the intestine, but not in hypodermis or muscle, reduced PLM axon degeneration in *mec-17(ok2109); lon-2(e678)* animals (Fig. 2G). Overexpressing *asah-1* in the mechanosensory neurons caused PLM axon outgrowth defects in *mec-17(ok2109); lon-2(e678)* animals (fig. S5), precluding analysis of PLM degeneration. Overexpression of *asah-1* in neurons of wild-type animals, either in the mechanosensory neurons (*mec-4* promoter) or pan-neuronally (*rab-3* promoter), also caused extensive axon outgrowth defects in the ALM and PLM neurons (Fig. 2H-I and fig. S5). However, no defects were detected in PVM guidance (fig. S5). These data reveal that neuronal *asah-1* overexpression disrupts axon outgrowth of the ALM and PLM axons, which extend axons embryonically, but not the PVM axons which extend post-embryonically. This potential embryonic effect is supported by the observation of axon guidance defects in the PVQ ventral nerve cord neurons when *asah-1* is overexpressed in the nervous system (fig. S5). However, animals overexpressing *asah-1* in the nervous system did not exhibit overt motility defects, suggesting that global nervous system architecture was intact. Together, these data reveal that intestinal *asah-1* induction reduces PLM axon degeneration in animals with defective microtubule stability, and that inappropriate neuronal expression of *asah-1*, with likely associated disruption of sphingolipid homeostasis, cell-autonomously causes neurodevelopmental defects.

Sphingolipids are amphipathic bioactive molecules with multiple cellular functions, including cell adhesion and migration, cell death and cell proliferation (*20*). Sphingolipid homeostasis is maintained through the *de novo* or salvage pathways (Fig. 3A) (*21*). Serine palmitoyltransferase (SPT) is the rate-limiting enzyme in the *de novo* pathway that generates ceramide - the ASAH-1 substrate (Fig. 3A). In *C. elegans, sptl-1* knockout causes embryonic lethality and larval arrest, likely due to loss of sphingolipid complexity. Therefore, to assess the importance of SPTL-1 in regulating PLM axon degeneration, we overexpressed *sptl-1* cDNA in the intestine. We found that *sptl-1* overexpression suppresses PLM axon degeneration of *mec-17(ok2109); lon-2(e678)* animals, confirming that ceramide or its derivatives are important for neuronal health (Fig. 3B). In the salvage pathway, lysosomal membrane ceramide is hydrolyzed to sphingosine by acid ceramidases (CDase), and sphingosine phosphorylation by sphingosine kinases (SphK) generates sphingosine-1-phosphate (S1P) (Fig. 3A) (*21*). We found that driving intestinal *sphk-1* expression suppresses PLM axon degeneration of *mec-17(ok2109); lon-2(e678)* animals, as we previously showed by overexpressing *sptl-1* or *asah-1* (Figs. 3B-D). Further, *sphk-1* loss increases PLM axon degeneration and prevents UA-induced reduction of PLM axon degeneration of *mec-17(ok2109); lon-2(e678)* animals (fig. S6). We have shown that intestinal overexpression of *asah-1* reduces PLM axon degeneration (Fig. 3C). To determine whether *sphk-1*, and therefore S1P generation, is required for ASAH-1 to perform this role we knocked down *sphk-1* expression in animals overexpressing *asah-1* in the intestine and evaluated PLM axon degeneration in *mec-17(ok2109); lon-2(e678)* animals. We found that *sphk-1* RNAi suppressed the beneficial effect of *asah-1* overexpression on PLM axon degeneration (Fig. 3E). Taken together, these data suggest that the protective role of UA in reducing PLM axon degeneration is dependent on S1P generation.

**Fig. 3.**
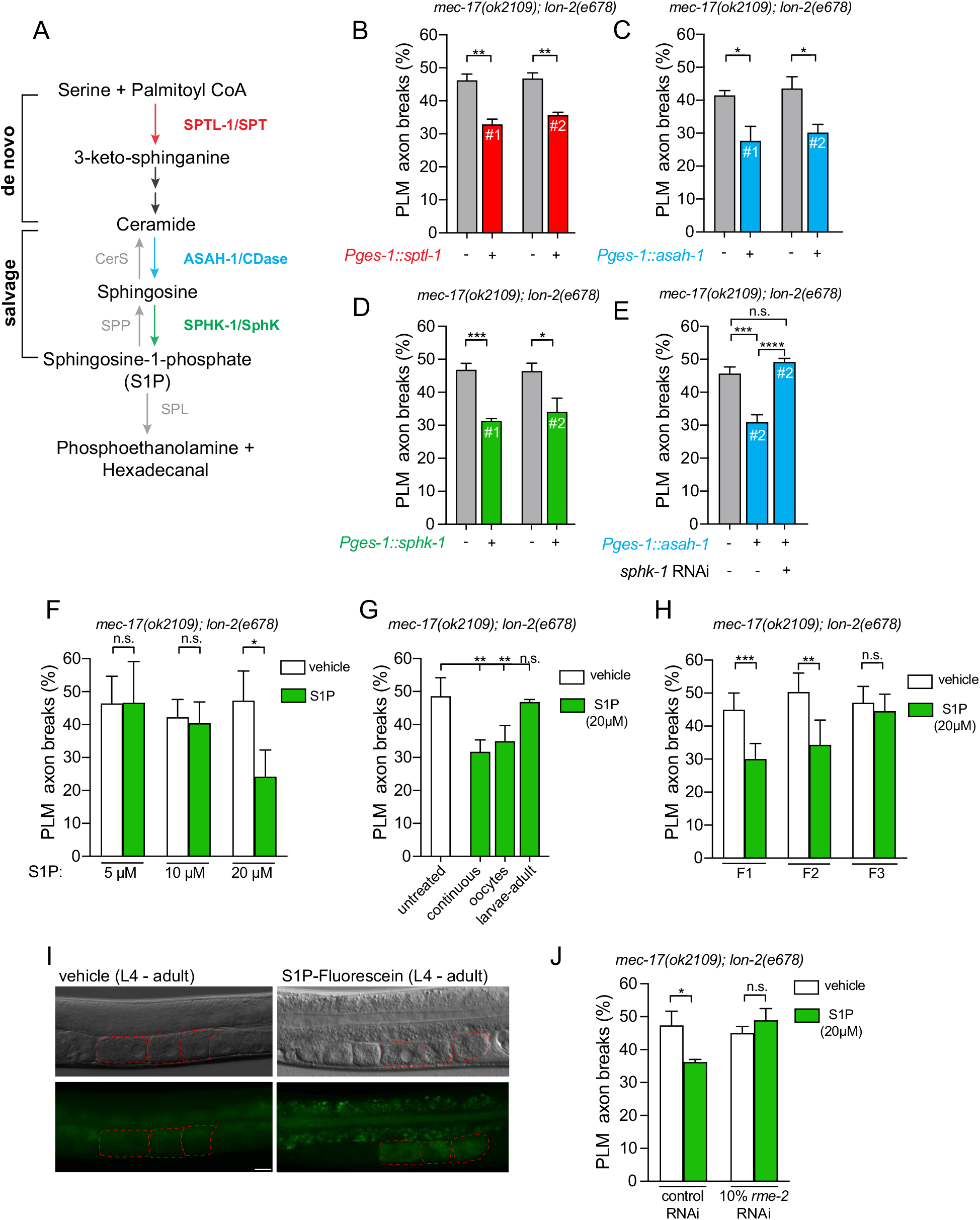
Sphingosine-1-phosphate intergenerationally protects the PLM neurons. (**A**) *C. elegans* orthologs of *de novo* and salvage sphingolipid pathway components. Sphingolipid metabolic enzymes (worm and mammalian ortholog) and sphingolipid intermediates are shown. The function of SPTL-1, ASAH-1 and SPHK-1 (colored lettering) in PLM axon degeneration were examined in (**B** to **E**). SPT, serine palmitoyltransferase; CerS, ceramide synthase; CDase, ceramidase; SphK, sphingosine kinase; SPP, sphingosine-1-phosphate phosphatase; SPL, sphingosine-1-phosphate lyase. (**B** to **D**) Driving *sptl-1* (**B**), *asah-1* (**C**) or *sphk-1* (**D**) cDNA in the intestine (*ges-1* promoter) rescues PLM axon breaks in *mec-17(ok2109); lon-2(e678)* animals. n = 70 to 80. # refers to independent transgenic lines. *P* values from one-way analysis of variance (ANOVA). \**P* ≤ 0.05; \*\**P* ≤ 0.01; \*\*\**P* ≤ 0.001; n.s., not significant. Error bars indicate SEM. (**E**) *sphk-1* RNAi knockdown suppresses the ability of intestinal *asah-1* expression to rescue PLM axon breaks in *mec-17(ok2109); lon-2(e678*) animals. n = 72 to 94. # refers to the transgenic line used (line 2 from **C**). *P* values from one-way analysis of variance (ANOVA). \*\*\**P* ≤ 0.001; \*\*\*\**P* ≤ 0.0001; n.s., not significant. Error bars indicate SEM. (**F**) Continuous exposure (P0, L4 larva to F1, adult) of sphingosine-1-phosphate (S1P) rescues PLM axon breaks in *mec-17(ok2109); lon-2(e678*) animals. n = 73 to 79. *P* values from one-way analysis of variance (ANOVA). \**P* ≤ 0.05; n.s., not significant. Error bars indicate SEM. (**G**) S1P exposure of P0 mothers for 16 h (L4 - young adult) but not F1 larvae rescues PLM axon breaks in *mec-17(ok2109); lon-2(e678*) animals. n = 104 to 135. *P* values from one-way analysis of variance (ANOVA). \*\**P* ≤ 0.01; n.s., not significant. Error bars indicate SEM. (**H**) S1P exposure in P0 mothers for 16 h (L4 - young adult) rescues PLM axon breaks in *mec-17(ok2109); lon-2(e678)* animals for two generations (F1 and F2). n = 153 to 163. *P* values from one-way analysis of variance (ANOVA). \*\**P* ≤ 0.01; \*\*\**P* ≤ 0.001; n.s., not significant. Error bars indicate SEM. (**I**) S1P-Fluorescein, a fluorescently labelled S1P analog, is detected in oocytes of wild-type adult hermaphrodites after 16 h of feeding. Vehicle-treated controls (left) and S1P-Fluorescein treatment (right). Nomarski micrograph (top) and fluorescent image (bottom) of the same animals. Hatched red lines = oocytes. Lateral view, anterior to the left. Scale bar: 25 μm. (**J**) *rme-2* RNAi knockdown suppresses the ability of S1P to rescue PLM axon breaks in *mec-17(ok2109); lon-2(e678)* animals. n = 104 to 107. *P* values from one-way analysis of variance (ANOVA). \**P* ≤ 0.05; n.s., not significant. Error bars indicate SEM.

To directly examine the role of S1P in PLM axonal health we incubated *mec-17(ok2109); lon-2(e678)* animals for one generation (P0 L4 - F1 3-day old adult) with different S1P concentrations. We found that 20 μM S1P reduced PLM axon degeneration (Fig. 3F). To determine the S1P functional period, we exposed *mec-17(ok2109); lon-2(e678)* animals to S1P from P0 L4 larvae-adult (oocytes present) or L1 larvae-adult (all larval stages and adult). S1P only reduced axon degeneration in F1 progeny when provided to hermaphrodites containing oocytes (P0 L4-adult) - the same functional period as UA (Fig. 3G compared to Fig. 1D). Additionally, incubating P0 animals from L4 larvae-adult (16 hours) with S1P reduced axonal degeneration for two generations - revealing an epigenetic intergenerational effect of this lipid (Fig. 3H). These data suggest that intestinal S1P is transported within the yolk to oocytes to promote PLM neuron health in subsequent generations. To examine whether S1P can undergo intestine-oocyte transport, we fed wild-type L4 larvae with heat-killed OP50 *Escherichia coli* containing 20 μM S1P-Fluorescein, a fluorescently labelled S1P analog (Fig. 3I). Fluorescence was observed in the intestinal tract within 1h of feeding suggesting that S1P-Fluorescein is not immediately metabolized (fig. S6). After 16h of S1P-Fluorescein feeding, we detected fluorescence in proximal oocytes, suggesting yolk-dependent intestine-oocyte transport (Fig. 3I). To determine the importance of intestine-oocyte S1P transport for preventing neurodegeneration we knocked down *rme-2* by RNAi in *mec-17(ok2109); lon-2(e678)* animals incubated with S1P. We found that S1P neuroprotection requires *rme-2* (Fig. 3J). These data reveal that S1P-dependent neuroprotection requires maternal information transfer from the intestine and oocyte through the low-density receptor RME-2 receptor.

How does UA induce *asah-1* expression in the intestine to enable intergenerational protection of the nervous system? As UA induces expression of the *asah-1* transcriptional reporter (Fig. 2E-F), we reasoned that transcription factors (TFs) related to intestinal stress or metabolic control may regulate *asah-1* expression. We examined the *asah-1* promoter for conserved TF binding motifs and surveyed publicly available ChIP-seq data for TFs exhibiting strong peaks upstream of the *asah-1* coding sequence (Fig. 4A and fig. S7). We identified ChIP-seq peaks for two TFs in the *asah-1* promoter: PQM-1, a GATA zinc-finger TF, and CEH-60/PBX, a TALE class (Three Amino Acid Loop Extension) TF (Fig. 4A and fig. S7). These ChIP-seq peaks coincide with putative binding sites we identified *in silico* (Fig. 4A and fig. S7). PQM-1 and CEH-60 function in the intestine to balance transcriptional networks that govern stress responses and nutrient supply to progeny (*22, 23*). We therefore investigated the potential role of these TFs in controlling PLM axon degeneration and *asah-1* expression.

**Fig. 4.**
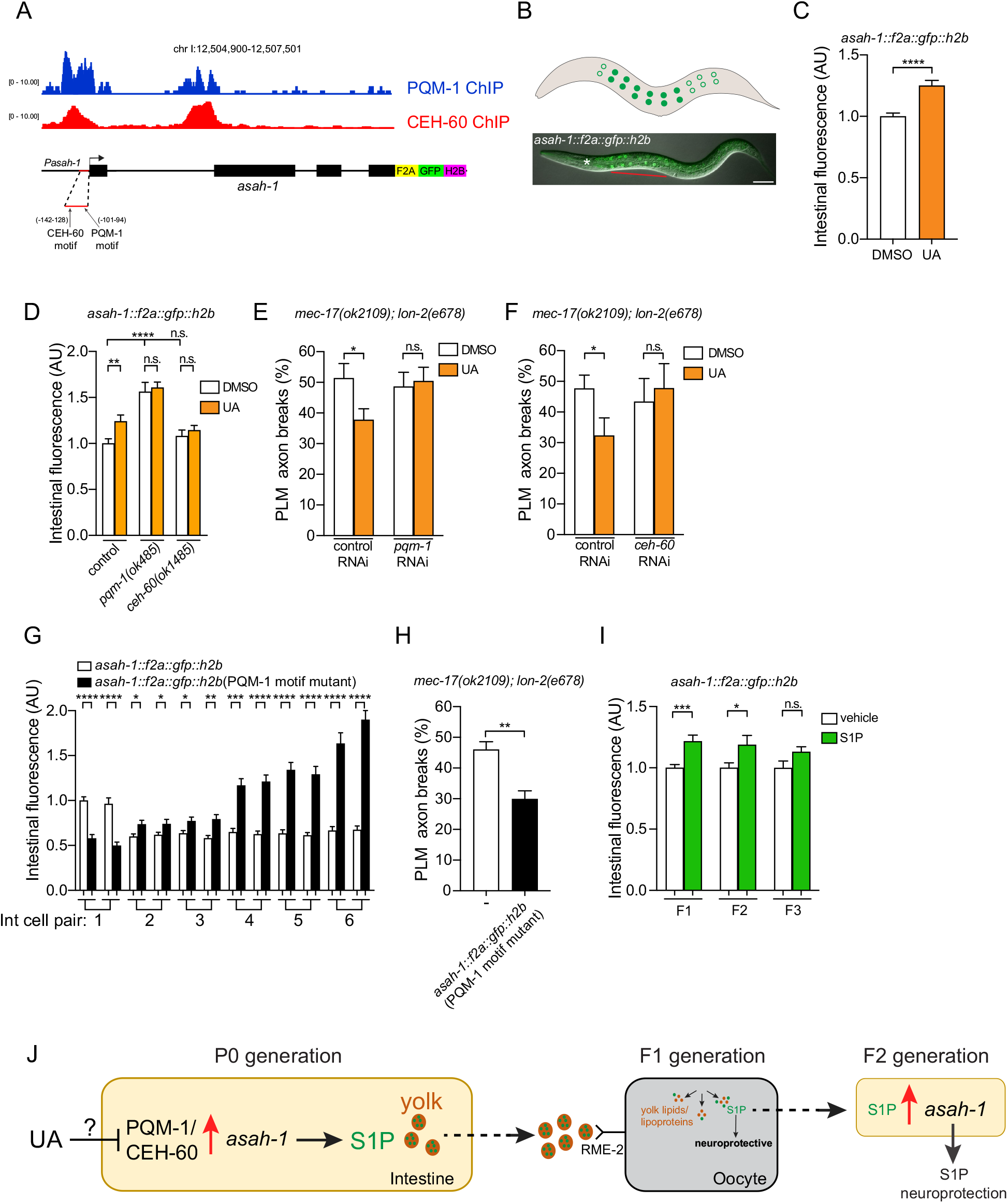
ASAH-1-mediated neuronal health promotion depends on PBX/MEIS-regulated transcription. (**A**) PQM-1 and CEH-60 ChIP-seq peaks at the *asah-1 cis*-regulatory region (top). Schematic of the CRISPR-Cas9-generated reporter of *asah-1* expression using F2A viral peptide-induced ribosomal skipping between endogenous *asah-1* and a *gfp-H2B* cassette. The *asah-1 cis-* regulatory region contains consensus binding sites for the CEH-60/PBX and PQM-1/C2H2-type zinc finger transcription factors (red line). Location of each motif (arrow) presented as distance from the ATG (+1). (**B**) GFP-H2B expression from the *asah-1::f2a::gfp::h2b* endogenous reporter (*rpls176*) is detected in the intestine (as with the transcriptional reporter, Fig. 2F.) from late embryogenesis (3-fold stage) through to adult (see fig. S7B for embryonic, early larval and adult images). Schematic (top) and overlay of Nomarski and fluorescence image of an L4 larva (bottom). GFP is weakly detected in the first pair and posterior intestinal cells (represented by open circles in the schematic). Lateral view, anterior to the left. Scale bar: 25 μm. (**C**) GFP-H2B expression from the *asah-1::f2a::gfp::h2b* endogenous reporter increases when animals are exposed to UA. Quantification of total nuclear fluorescence from 6 pairs of intestinal cells per worm is shown (see fig. S8A for individual intestinal cell measurements). n = 28 to 30. *P* value assessed by unpaired t test. *\*\*\*\*P* ≤ 0.0001. Error bars indicate SEM. (**D**) *pqm-1(ok485)* and *ceh-60(ok1485*) mutants prevent UA induction of the *asah-1::f2a::gfp::h2b* endogenous reporter. Quantification of total nuclear fluorescence from 6 pairs of intestinal cells per worm is shown. n = 18 to 26. *P* value assessed by unpaired t test. \*\**P* ≤ 0.01, \*\*\*\**P* ≤ 0.0001. Error bars indicate SEM. (**E** and **F**) *pqm-1* or *ceh-60* RNAi knockdown suppresses UA-induced rescue PLM axon breaks in *mec-17(ok2109); lon-2(e678*) animals. n = 74 to 103 (**E**) and 65 to 151 (**F**). *P* values from one-way analysis of variance (ANOVA). *\*P* ≤ 0.05; n.s., not significant. Error bars indicate SEM. (**G**) Mutation of putative PQM-1 binding motif in the *asah-1* promoter induces GFP-H2B expression from the *asah-1* endogenous reporter. Quantification of nuclear fluorescence from each pair of intestinal cells per worm is shown (Int cell pair 1 = second pair of intestinal cells from the pharynx). n = 30 to 32. *P* value assessed by unpaired t test. \**P* ≤ 0.05, \*\**P* ≤ 0.01, \*\*\**P* ≤ 0.001, \*\*\*\**P* ≤ 0.0001. Error bars indicate SEM. (**H**) Mutation of the putative PQM-1 binding motif in the *asah-1* promoter reduces PLM axon breaks in *mec-17(ok2109); lon-2(e678)* animals. n = 100 to 104. *P* values from one-way analysis of variance (ANOVA). \*\**P* ≤ 0.01. Error bars indicate SEM. (**I**) S1P exposure in P0 mothers for 16 h (L4 - young adult) increases GFP-H2B expression from the *asah-1::f2a::gfp::h2b* endogenous reporter for two generations (F1 and F2). n = 23 to 28. *P* values from one-way analysis of variance (ANOVA). \**P* ≤ 0.05; \*\*\**P* ≤ 0.001; n.s., not significant. Error bars indicate SEM. (**J**) Model summarizing how UA and S1P intergenerationally protect the nervous system.

To directly assess whether PQM-1/CEH-60 transcriptionally regulate endogenous *asah-1*, we generated a fluorescent reporter by inserting a *f2A-gfp-h2B* cassette immediately downstream of the *asah-1* coding sequence using CRISPR-Cas9 (Fig. 4A-B and fig. S7). Ribosomal skipping occurs at the viral *f2A* sequence, and thus independently translated GFP-H2B protein is visualized in nuclei (Fig. 4B and fig. S7) (*24*). We detected GFP-H2B expression in anterior intestinal nuclei (except the most anterior nuclei pair), with weak/undetectable expression in the posterior intestine (Fig. 4B and fig S7). This anteriorly biased expression was also observed in the *asah-1* transcriptional reporter (Fig. 4B compared to fig. S4), suggesting that spatial expression of *asah-1* is transcriptionally regulated. Exposing the *asah-1* endogenous reporter to UA induced GFP-H2B expression (Fig. 4C and fig. S8A), supporting our previous finding with the *asah-1* transcriptional reporter (Fig. 2F). We crossed *pqm-1(ok485)* and *ceh-60(ok1485)* loss-of-function mutants into *asah-1::f2A-gfp-h2B* animals and measured GFP-H2B fluorescence in intestinal nuclei. Loss of PQM-1 elevated GFP-H2B expression and loss of either PQM-1 or CEH-60 prevented GFP-H2B induction by UA (Fig. 4D). To examine the potential role of PQM-1 and CEH-60 UA-induced neuroprotection, we knocked down their expression using RNAi (Fig. 4E-F). *ceh-60* and*pqm-7* RNAi inhibited UA neuroprotection in *mec-17(ok2109); lon-2(e678)* animals (Fig. 4E-F). Hence, PQM-1 and CEH-60 enable *asah-1* induction and neuroprotection in response to UA exposure.

As PQM-1 and CEH-60 have key roles of in controlling metabolic homeostasis and yolk production (*22, 23*), *asah-1* regulation in these TF mutants may be secondary consequence of disrupted lipid homeostasis. Therefore, to directly examine the importance of PQM-1 and CEH-60 binding on *asah-1* expression, we used CRISPR-Cas9 to mutate putative binding motifs that coincide with the PQM-1/CEH-60 ChIP peaks in the *asah-1* promoter (Fig. 4A and figs. S7). We found that mutating the PQM-1 binding motif strongly induced GFP-H2B intestinal expression, with mutation of the CEH-60 site having a weaker effect (Fig. 4G and fig. S8B). These data support a direct role for these TFs in *asah-1* transcriptional repression and suggests that control of PQM-1/CEH-60 occupation at the *asah-1* promoter can control sphingolipid homeostasis. We also noticed that mutating the PQM-1 motif caused a posterior shift in GFP-H2B expression, revealing spatial regulation of intestinal expression (Fig. 4G). As mutating the PQM-1 motif strongly increased expression from the *asah-1* locus, we asked whether this was neuroprotective. When the *asah-1::f2A-gfp-h2B* (PQM-1 motif mutant) strain was crossed into *mec-17(ok2109); lon-2(e678*) animals, we observed robust reduction in PLM axon degeneration (Fig. 4H). This reveals that transcriptional de-repression of *asah-1* has a functionally relevant outcome for neuronal health.

Ceramide hydrolysis is the only catabolic source of sphingosine, and therefore ceramidase (e.g. ASAH-1) activity is deemed a rate-limiting step in governing intracellular sphingosine and S1P levels (*25*). We hypothesized that S1P intergenerational inheritance may be triggered by an imbalance in sphingolipid homeostasis, such as increased S1P levels. Thus, detection and recycling of S1P to sphingosine and then ceramide may feedback to induce *asah-1* expression in progeny. To explore this possibility, we exposed *asah-1::f2A-gfp-h2B* P0 animals to S1P for 16 h (L4 - young adult) and measured GFP-H2B intestinal expression in subsequent generations. Remarkably, GFP-H2B levels were increased in F1 and F2 progeny (Fig. 4I), establishing a regulatory mechanism for S1P-induced intergenerational inheritance.

Together, our findings show that neuronal development and health require spatial regulation of sphingolipid biosynthetic enzymes. Elevated acid ceramidase expression in the intestine is neuroprotective, whereas neuronal expression causes axon outgrowth developmental defects. The neuroprotective effect of intestinal acid ceramidase is dependent on a sphingosine kinase that phosphorylates sphingosine to generate S1P. In support of this, maternally derived S1P is transferred from the intestine to oocytes to protect against axon degeneration (Fig. 4J). Maternal S1P is likely transported within low-density yolk lipoproteins, as removal of the RME-2 yolk receptor inhibits S1P neuroprotection. In mammals, plasma lipoproteins carry S1P, highlighting a conserved mode of carriage (*26*). We discovered that elevated S1P in mothers intergenerationally increases expression of ASAH-1 - the rate-limiting enzyme in the sphingolipid salvage pathway. This finding may suggest a broader underlying role for metabolism and metabolic gene expression in intergenerational effects across species. Our data show that maternal S1P reduces neurodegeneration in axons with impaired cytoskeletal function. We propose that elevated S1P in the oocyte/embryo influences the axonal environment to limit axon fragility. The important roles of sphingolipid homeostasis in plasma membrane fluidity, lipid raft integrity and the molecular trafficking may be central to the protective role S1P plays in this context.

## Acknowledgments

We thank Brent Neumann and members of the Pocock laboratory for comments on the manuscript. Some strains used in this study were provided by the *Caenorhabditis* Genetics Center, which is funded by NIH Office of Research and Infrastructure Programs (P40 OD10440) and the National BioResource Project (NBRP) Japan.

## Funding

National Health and Medical Research Council grants GNT1105374, GNT1137645 and GNT2000766 (RP)

Apex Biotech Research Pty Ltd donation 281589068 (ZX)

## Author contributions

Conceptualization: WW, ZX, RP

Methodology: WW, JL, RP

Investigation: WW, TS, XC, QF, RC, RP

Visualization: WW, XC, RC, RP

Funding acquisition: ZX, RP

Project administration: WW, RP

Supervision: JL, ZX, RP

Writing – original draft: RP

Writing – review & editing: WW, TS, XC, QF, RC, JL, ZX, RP

## Data and materials availability

All data are available in the main text or the supplementary materials.

## Supplementary Materials

Materials and Methods

Supplementary Text

Figs. S1 to S8

Data S1 to S5

