## Supplementary figures and images for "An intestinal sphingolipid promotes neuronal health across generations"

### Fig S1

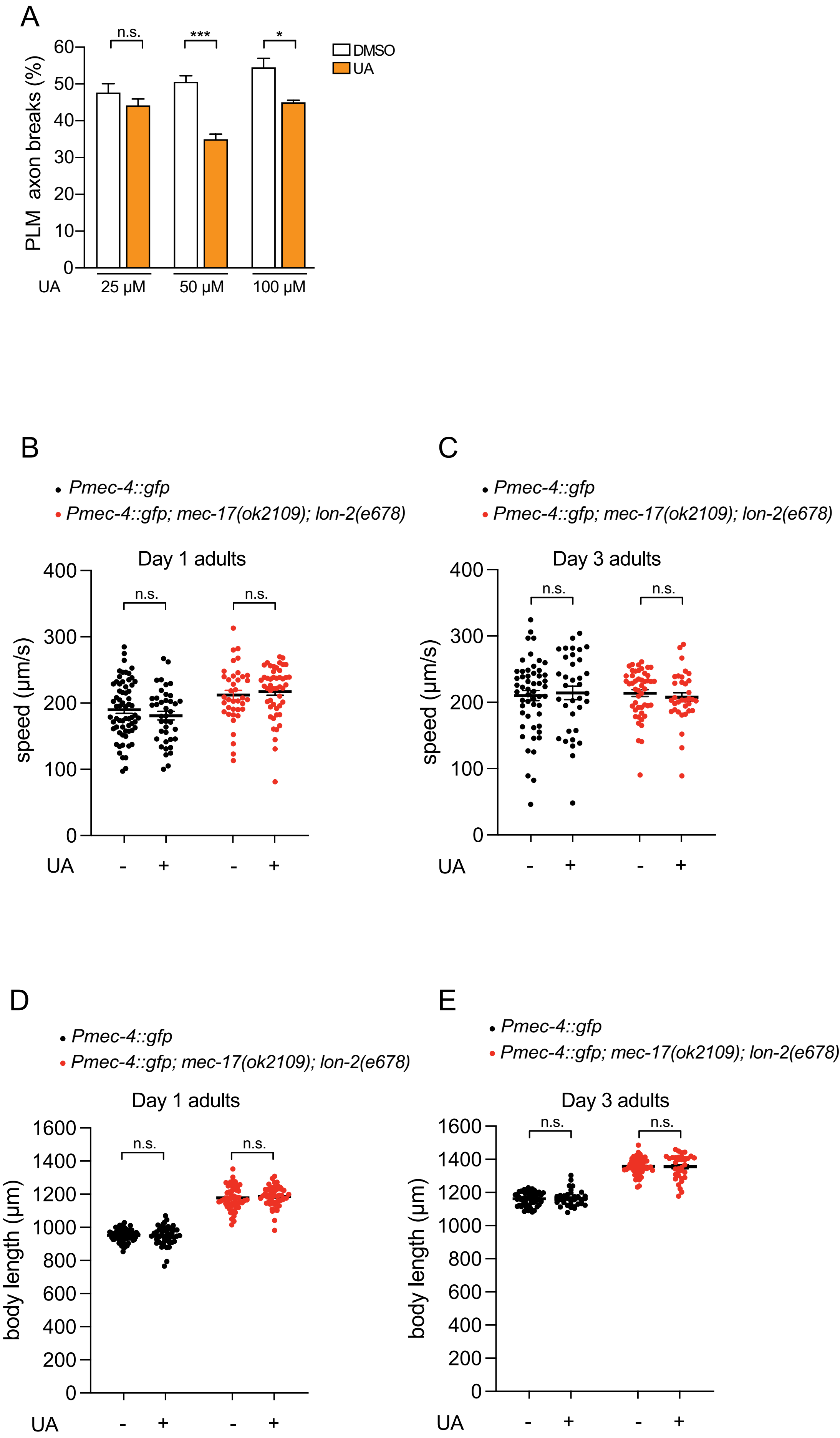

### Fig S2

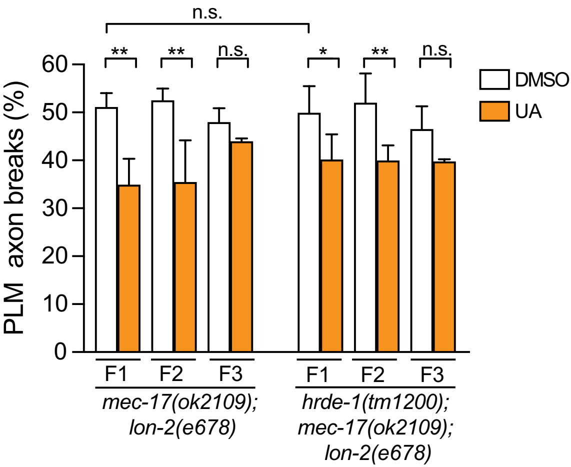

### Fig S3

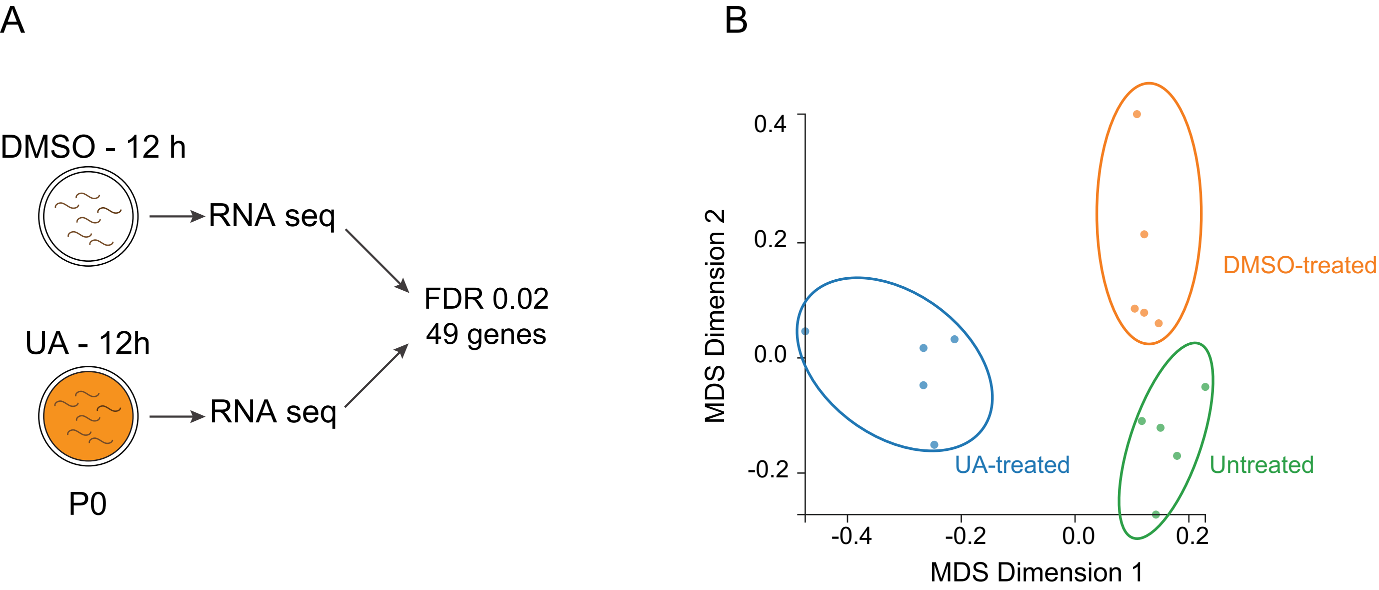

### Fig S4

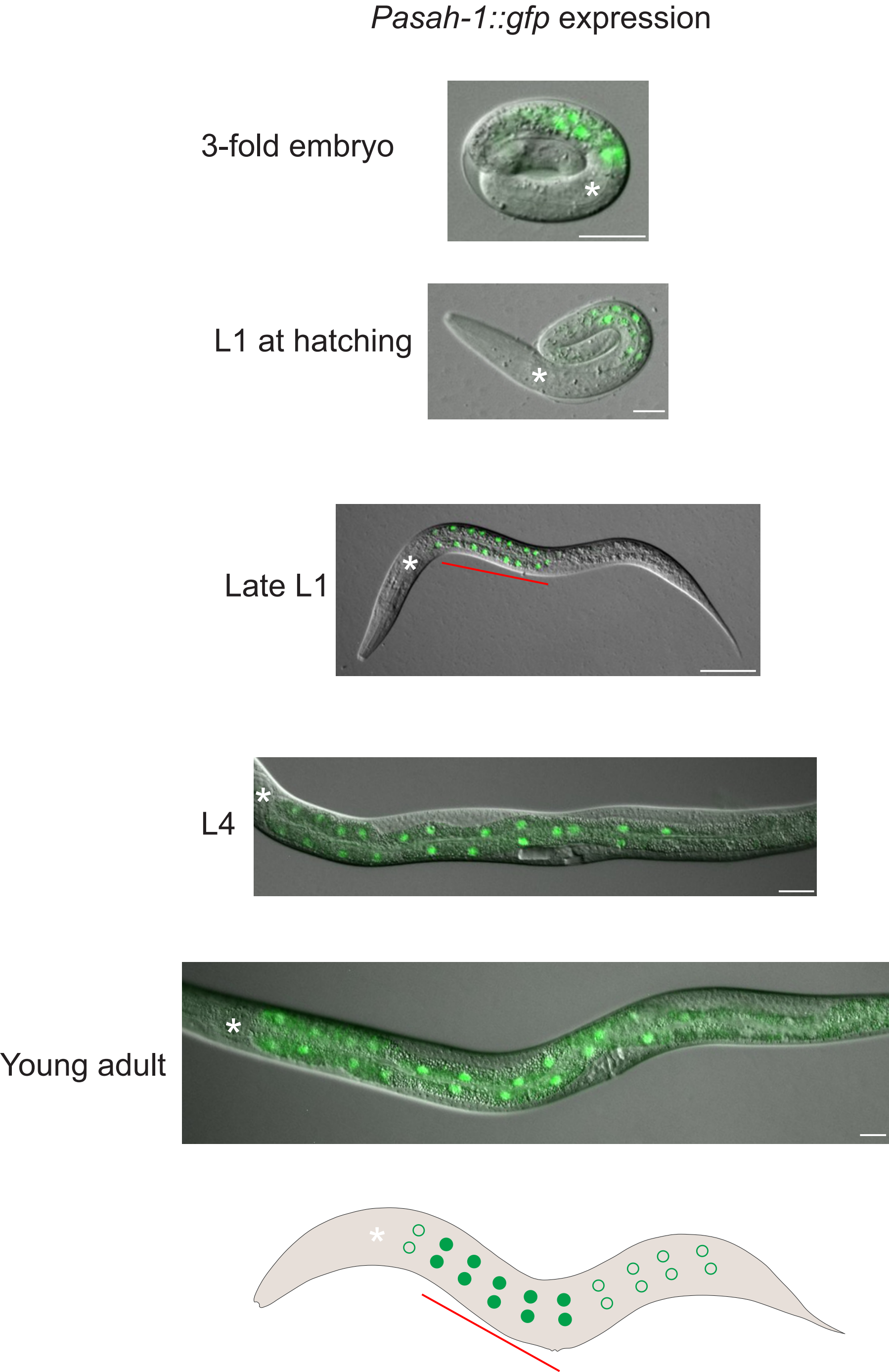

### Fig S5

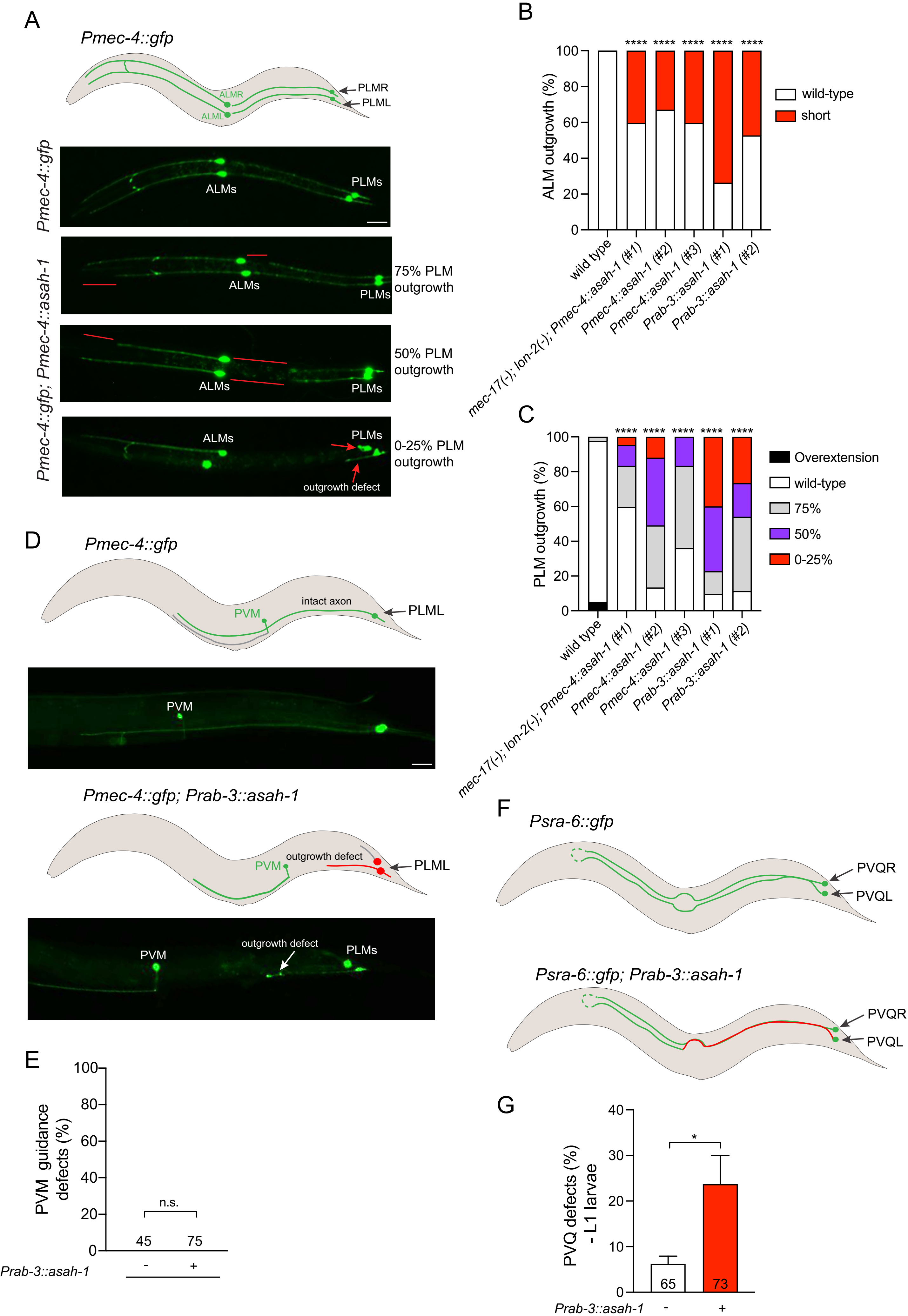

### Fig S6

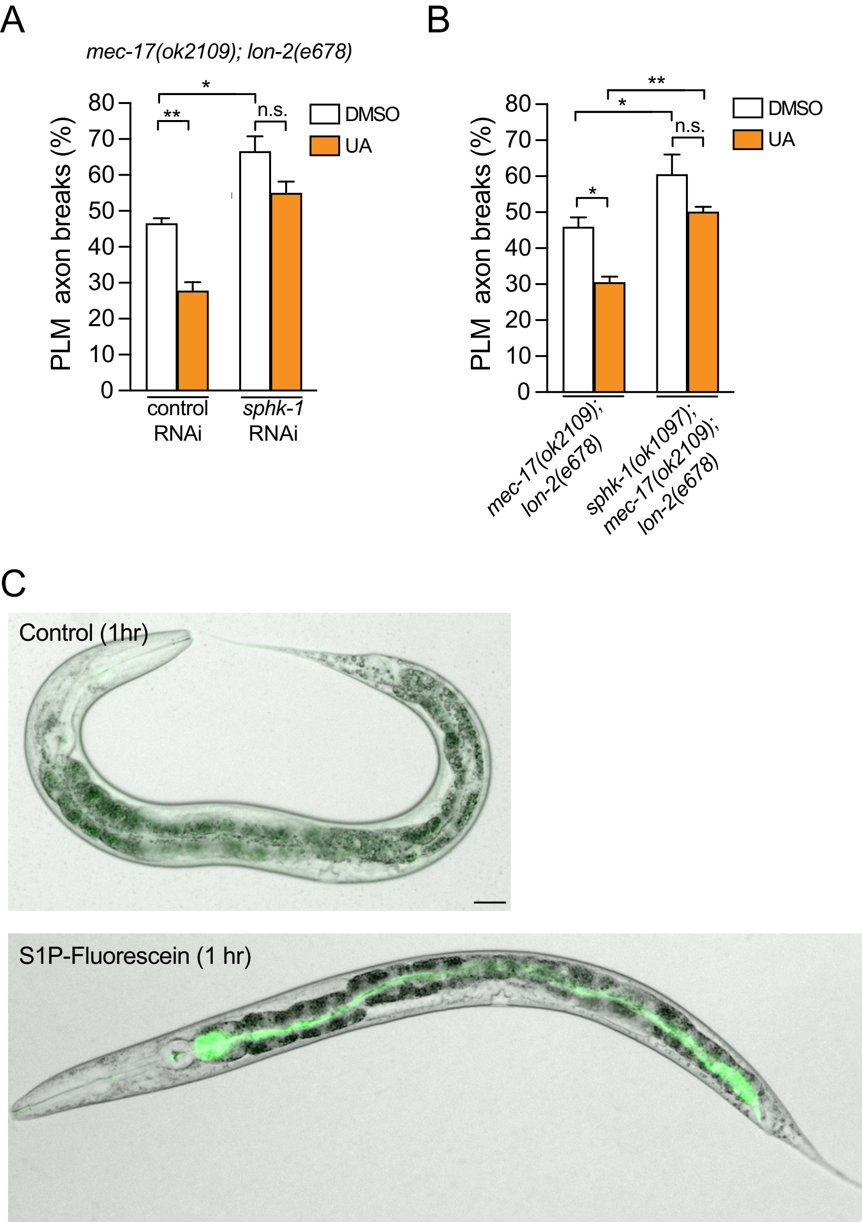

### Fig S8

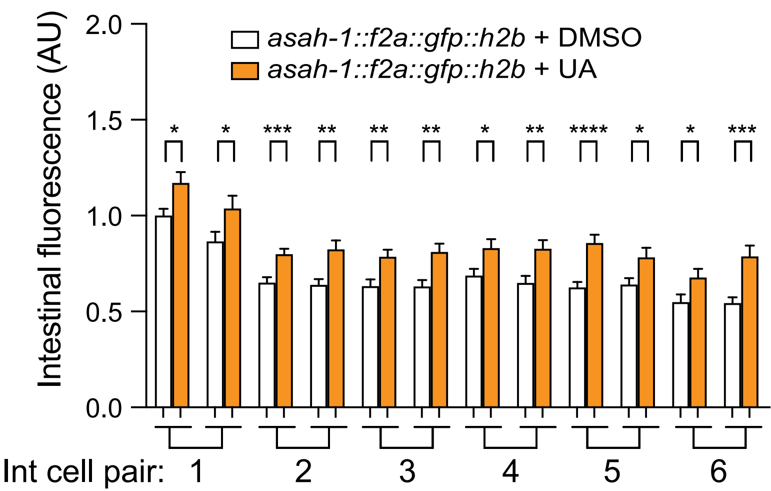

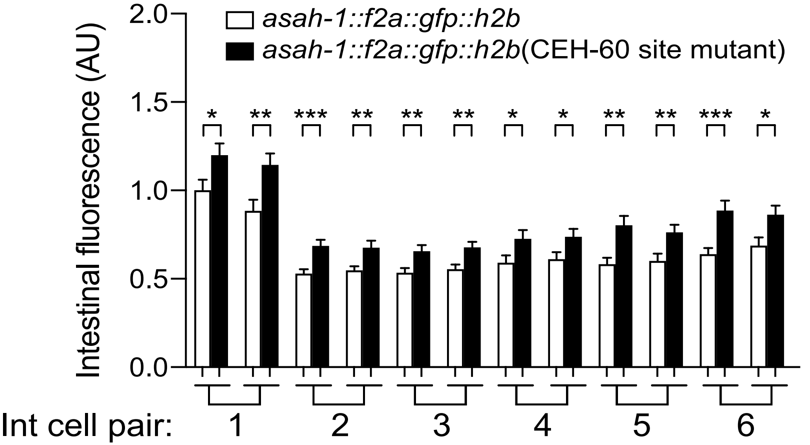
