## Supplementary material for "An intestinal sphingolipid promotes neuronal health across generations": Fig S7

A

***asah-1* promoter (200bp)**

TTTAATGAATTCTTCTCTGAAAACGCTTGTAAATTATGCTCCATAAACCAAAAACC
GGTGTGATTGATGACATTTTCATGAGGGAAATAAATTAAAAAGTGATAAGGAGTA
AAGTGTGGCCACGTGTTTTCCGCAAAAAGTATTGATAAGTCGCCTCGCGGAGC
ACACGCTTTGCCACTATTCAGAGCTAGTCAAGAAAG

TGTGATTGATGACA = CEH-60 motif
TGATAAG = PQM-1 motif #1

**Mutations introduced to the *asah-1* promoter using CRISPR-Cas9**

TTTAATGAATTCTTCTCTGAAAACGCTTGTAAATTATGCTCCATAAACCAAAAACC
GGTGCGGCCGCTGACATTTTCATGAGGGAAATAAATTAAAAAGTGGGATCCGA
GTAAAGTGTGGCCACGTGTTTTCCGCAAAAAGTATTCCATGGTCGCCTCGCGG
AGCACACGCTTTGCCACTATTCAGAGCTAGTCAAGAAAG

B

*asah-1::f2a::gfp::h2b* 3-fold embryo

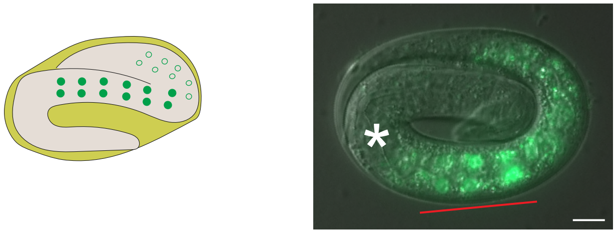

*asah-1::f2a::gfp::h2b* L1 larva

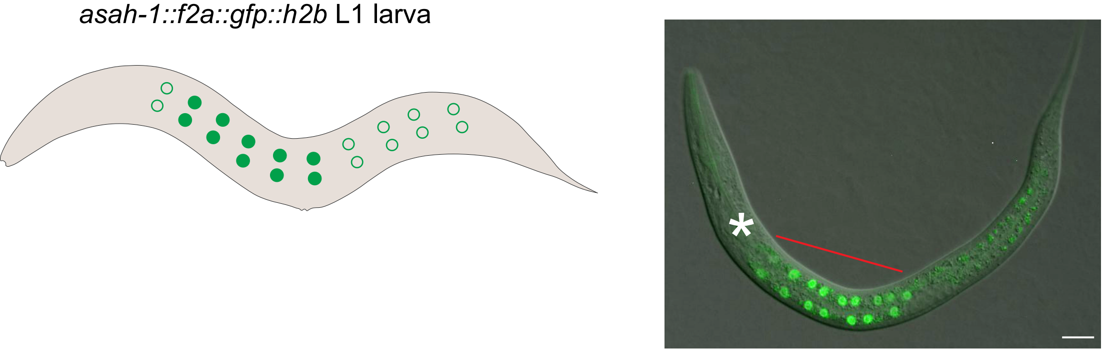
